# How compromised is reproductive performance in the endangered North Atlantic right whale? A proposed method for quantification and monitoring

**DOI:** 10.1101/2023.11.21.568115

**Authors:** Timothy R. Frasier, Philip K. Hamilton, Richard M. Pace

## Abstract

The endangered North Atlantic right whale (*Eubalaena glacialis*) showed limited recovery from the cessation of industrial whaling until 2011, and has since been in decline. Research is therefore focused on identifying what factors are limiting recovery and what conservation actions will be most effective. A compromised reproductive rate is one of the primary reasons for this lack of recovery, yet there is no consensus on how to quantify reproductive performance. As one potential solution, we propose a relatively simple approach where we calculate the theoretical maximum number of calves each year. Comparing this expected number to those observed provides a means to quantify the degree to which reproduction is being compromised and trends thereof over time. Implementing this approach shows that, between 1990 and 2017, the number of calves born never came close to the theoretical maximum, resulting in overall reproductive performance being only about 27% of that expected. In addition to quantifying the magnitude of the reproductive problem, this approach should also be useful for quantifying the role of reduced reproductive performance in limiting species recovery, and for aiding research programs focused on identifying what factors are compromising reproduction.

## 1 INTRODUCTION

There is increasing concern regarding the viability of the endangered North Atlantic right whale (*Eubalaena glacialis*). This species was once the target of intensive whaling (Reeves et al. 2007), but has not recovered well despite the cessation of that directed persecution. Specifically, the species showed slight, yet heterogeneous, signs of increase from the initiation of dedicated studies in 1980 through about 2011. The rate of this increase was ∼2.5%/year (Knowlton et al. 1994; Pace et al. 2017), which was substantially slower than the recovery rates of several right whale populations in the southern hemisphere (representing a different species: *E. australis*), which are increasing at rates of about 7%/year (Bannister, 2001; Cooke et al. 2001; Best et al. 2001). However, recovery in the North Atlantic was slow and heterogeneous enough as to be all but imperceptible at the time (Best, 1993; Clapham et al. 1999), and only became unequivocal in hindsight from a long-term perspective decades later (Pace et al. 2017). In addition to this reduced rate of recovery for decades, the species has been in decline since 2011 (Pettis et al. 2021).

This lack of recovery has been attributed to two major factors: a high rate of anthropogenic mortality from vessel strikes and entanglement in fishing gear (Knowlton et al. 2012; Pace et al. 2021), and a reproductive rate that is markedly lower than their known potential (Kraus et al. 2001; Kraus et al. 2007; Browning et al. 2010). While many conservation efforts have been directed at lowering rates of anthropogenic mortality (e.g., Vanderlaan et al. 2008; van der Hoop et al. 2015; Moore 2019), the factors possibly compromising reproductive performance are not yet well-understood and are likely multifaceted (Kraus et al. 2007). To date, evidence suggests that nutritional stress and sublethal effects of entanglements are at least partially responsible (Meyer-Gutbrod et al. 2015; Stewart et al. 2021, 2022), but other factors are also likely contributing (e.g., Frasier et al. 2015).

Demographic descriptors of wildlife populations often include measures of reproduction and survival rates. These can be expressed solely as rates (*per capita*) or as absolutes (total natality and total mortality). For North Atlantic right whales, high quality measurements exist for these metrics with respect to *observed* reproductive rates, but there are not currently any clear benchmarks of *expected* reproductive potential, against which observed values can be compared to better quantify the magnitude and trends of the reproductive problem and the degree to which it is limiting recovery.

Our goal was to develop a simple and clear method with which to gauge observed reproductive output against a theoretical benchmark based on the reproductive cycle and age structure of the North Atlantic right whale population. The hope is that such quantification will be helpful in quantifying the reproductive problem and the degree to which reproduction is limiting species recovery. Doing so should also as aid research focused on identifying what factors are limiting reproductive performance by quantifying the magnitude and trends of the problem itself.

## 2 METHODS

### 2.1 Data

Analyses were based on the reproductive histories of females from 1990-2017. Briefly, individual North Atlantic right whales can be identified based on their distinguishing physical characteristics: primarily callosity patterns on their heads, but also including variation in skin pigmentation and scarring (Payne et al. 1983; Kraus et al. 1986a). Dedicated field studies began in 1980, and fieldwork now takes place almost year-round and throughout the range of the species (Brown et al. 2007). The ability to recognize individuals across space and time has provided a wealth of individual-based information (Hamilton et al. 2007). Moreover, there is one identified calving area for the species, which has been intensively surveyed since 1990, resulting in thorough reproductive histories for each female (Kraus et al. 1986b; Brown et al. 2007). Data on female age and reproductive histories were obtained from the North Atlantic Right Whale Consortium (www.narwc.org) – a collaborative group of individuals and entities involved in right whale research and conservation, which also serves as a means for data sharing.

### 2.2 Characterization of the reproductive performance

Prior to developing a method to quantify the reproductive performance, it was first necessary to assess the contributing components to ensure that they are adequately captured with the new method. Two major components of poor reproductive performance have been identified: (1) those associated with inter-birth intervals; and (2) females that delay reproduction until much later in life than average, or that are completely nulliparous.

For inter-birth intervals, North Atlantic right whale females are capable of giving birth once every three years (Knowlton et al. 1994; Kraus et al. 2001; Kraus et al. 2007). This represents approximately one year of lactation, one year of “resting” to replenish tissue and energy stores, and then one year of gestation. However, there is a great deal of variation around this 3-year interval at temporal and individual scales. Temporally, the average inter-birth interval for calving females fluctuates over time, and has increased during periods where the species appeared to be nutritionally stressed (Kraus et al. 2001; Miller et al. 2011; Moore et al. 2021). Second, there is a high degree of variability in the average inter-birth interval across different females: where some females reproduce fairly regularly at 3-or 4-year intervals, whereas other females have much longer average intervals. Occasionally, upon losing a calf prior to or during migration to northern feeding grounds (early in her lactation period), females may produce a calf after only 2 years (Knowlton et al. 1994; Payne et al. 1990).

The second major component of reduced reproductive performance involves increased age of primaparity, and females that never reproduce and are therefore nulliparous. Quantifying the number of truly nulliparous females is difficult due to the long lifespan of right whales (unknown, but estimated at >70 years, Hamilton et al. 1998; Katona & Kraus 1999), making it unclear if some of the females may have reproduced prior to the beginning of study in 1980, and just not since then. Regardless, it is currently estimated that about 12% of adult females are truly nulliparous (Kraus et al. 2007). Moreover, although the average age of first reproduction is approximately 8 years for females, some do not calve until much older, with a recent assessment indicating that half of the females between the ages of 10 and 19 have not yet given birth (Moore et al. 2021; Reed et al. 2022).

To assess the relative contribution of both components on reproductive patterns, we characterized the reproductive histories of adult females in each year. This involved identifying how many adult females were “presumed alive” in each year (the majority of mortalities are not detected (Pace et al. 2021), and therefore individuals are “presumed alive” until their sixth year without being sighted (Knowlton et al. 1994; Hamilton et al. 2007)), and what their reproductive histories were up to and including that year: (1) those that had not reproduced yet; (2) those that had produced one calf; and (3) those that had reproduced two or more times, and were therefore available for analyses of inter-birth intervals. This resulted in a “reproductive cross-section” of the adult females in each year, with respect to these categories.

### 2.3 Quantifying reproductive performance

In the past, consideration of the reproductive problem has largely focused on inter-birth intervals and variations thereof over time (e.g., Kraus et al. 2001, Pettis et al. 2021). However, doing so does not include nonreproductive females and females that only calve once. It was therefore desirable to develop a method that considers both inter-birth intervals as well as the proportion of females that are reproducing.

Our proposed approach is based on calculating the *expected* number of calves born in each year, and using that as a metric against which the actual number of calves born is compared. We argue that the expected number of calves born can be calculated as the number of living adult females minus those that gave birth in the previous two years. The rationale is that if reproduction was not being compromised then we would expect each adult female to give birth approximately once every three years. This approach captures both manifestations of the reproductive problem. First, females and/or time periods with longer inter-birth intervals will result in fewer calves born than expected because fewer females will be reproducing on the expected three-year schedule. Second, non-reproducing females will also cause the observed number of calves to be lower than expected because they are not producing calves.

Because this approach hinges on the logic that if reproduction was not being compromised by one or more factors, then females would be giving birth once every three years, it is worthwhile to justify this perspective. First, and perhaps most convincingly, we know that North Atlantic right whale females can reproduce once every three years because some of them have before (Knowlton et al. 1994; Kraus et al. 2001; Pettis et al. 2021). Indeed, the average inter-birth interval from 1980 through 1992 was 3.67 years (Knowlton et al. 1994). Thus, it is clearly physiologically possible for them to do so under favorable conditions. Second, the average inter-birth interval for females in many populations of the closely related southern right whale (*E. australis*) is approximately three years (South Africa – 3.12 years, Best et al. 2001; Argentina – 3.35 years, Cooke et al. 2001; Australia – 3.33 years, Burnell 2001). Lastly, data from many other baleen whale species indicate that the proportion of reproducing females is closely aligned with known calving intervals. For example, the average inter-birth interval for female humpback whales (*Megaptera novaeangliae*) in the North Atlantic is ∼2.4 years, and the estimated percentage of adult females that are pregnant each year closely matches this at 41% (Clapham & Mayo 1988). Many other studies of marine mammals and other large mammals show a close relationship between inter-birth intervals and pregnancy rates (e.g., Hammill & Gosselin 1995; Herzing 1997; Festa-Bianchet et al. 1998; Clutton-Brock et al. 2003), indicating that, in a healthy population, the vast majority of females reproduce at their maximum physiological rate (at least for species that do not live in social groups where the reproductive success of some females is controlled by others).

Another aspect of reproductive biology to consider is senescence: if North Atlantic right whale females go through a period of senescence then it would not be appropriate to include them in calculations of the expected number of calves in each year. However, although senescence is clearly evident in some toothed whale species (Marsh & Kasuya 1986; Ellis et al. 2018), life-history theory does not predict a period of senescence in baleen whales, nor has evidence of it been found, despite a number of long-term individual-based studies of many baleen whale species and populations (Marsh & Kasuya 1986; Whitehead & Mann 2000).

Combined, these data suggest that if reproduction was not being compromised, it would be reasonable to expect North Atlantic right whale females to reproduce approximately once every three years. Based on this logic, we calculated the expected number of calves to be born in each years as the number of adult females alive in that year minus the number of females that gave birth within the previous two years. These expected number of calves born were calculated as the median number of females estimated to be adults and alive in each year who did not reproduce in the previous two years, from the model described in Pace et al. (2017), which also provided estimates of the uncertainty around these numbers for each year.

## 3 RESULTS

The reproductive histories of adult females presumed alive in years 1990-2017 are shown in **Figure 1A**, and a cross-section of those females for the year 2017 is shown in **Figure 1B**. The data show a clear decreasing trend in females that have reproduced multiple times, with a corresponding increase in the number of females that have not yet reproduced or have had just one calf. In 2017 there were 134 females presumed to be alive and adult. Of these 134 females, 75 (56%) had given birth to more than one calf and were therefore available for analyses of inter-birth intervals. Twenty-eight of these females (21%) had given birth to just one calf (as of 2017). The remaining 31 individuals (23%) had not yet given birth.

**Figure 1.**
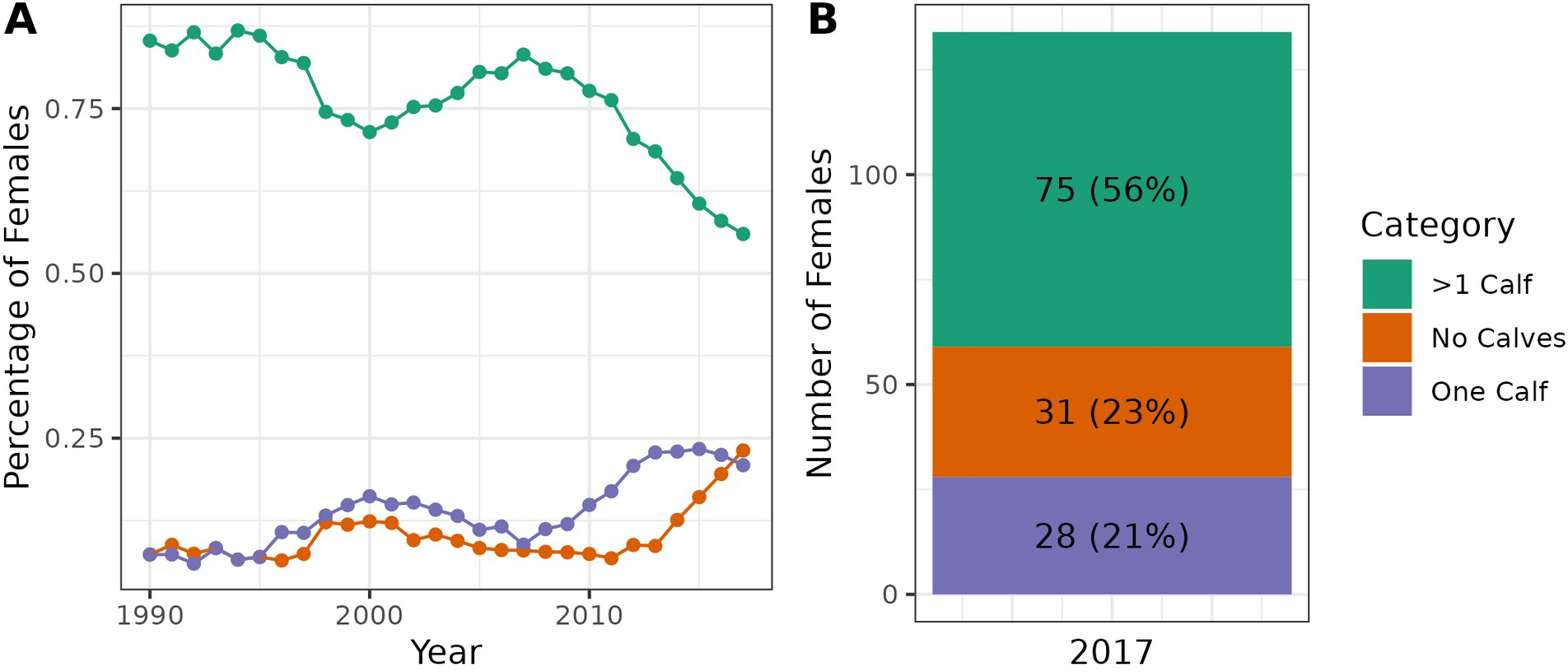
Reproductive histories of adult females presumed alive in years 1990-2017. Females were binned in one of three categories for each year: (1) no calves as of that year (i.e., they had not successfully reproduced yet), (2) one calf as of that year, or (3) having given birth to >1 calf as of that year. Only females that have given birth to multiple calves (i.e., those in the “>1 Calf” category) can be used for analyses of inter-birth intervals. **(A)** The percentage of adult females that were in each category across years. **(B)** A “cross-section” of the adult females alive in the year 2017.

The expected number of calves born for the years 1990-2017, and associated 95% highest density intervals (HDIs), are shown in **Figure 2A**. Also shown are the observed number of calves born in each year. **Figure 2B** shows the same data in a different manner, where the observed number of calves are plotted as a percentage of those that were expected for each year. Although there is substantial annual variability, the average over time shows that the reproductive performance of this species is slightly less than a third (∼27%) of that expected.

**Figure 2.**
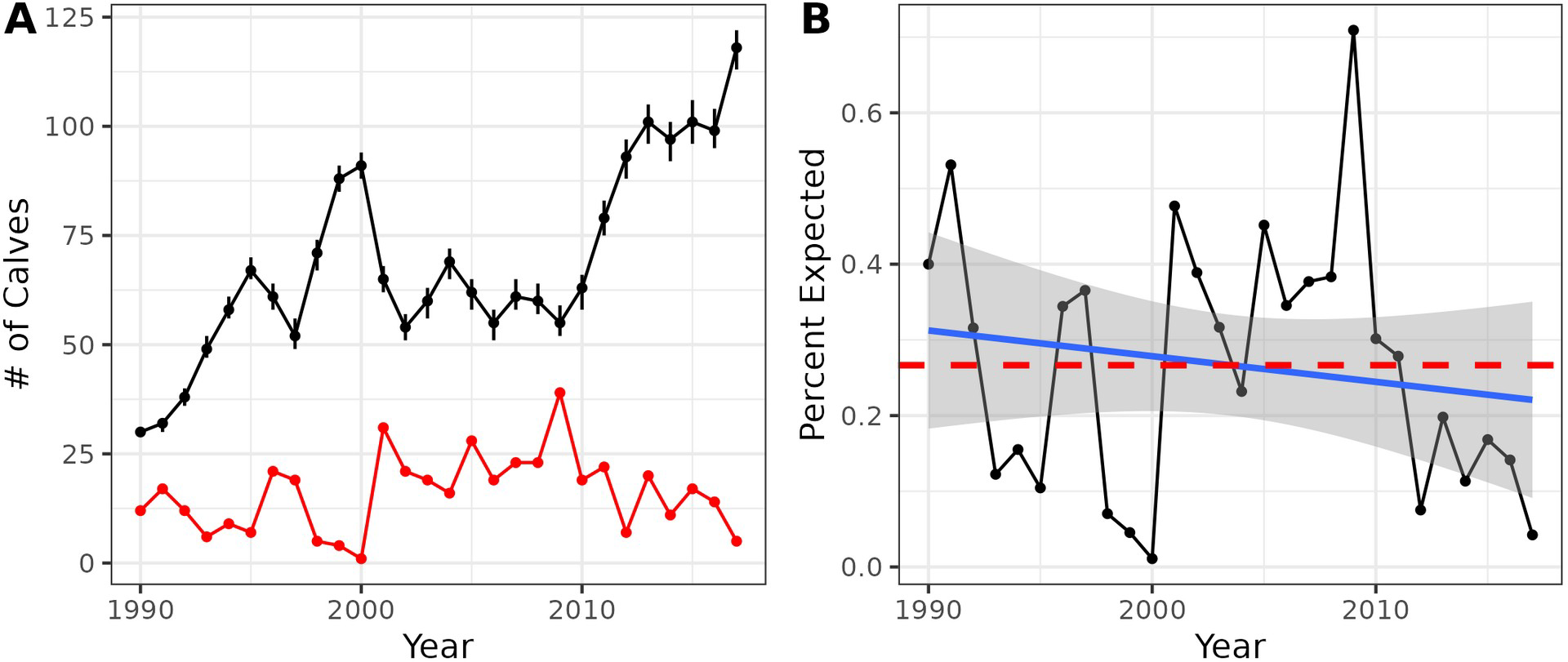
Expected and observed number of calves born in each year from 1990-2017. **(A)** The expected number of calves born each year (black circles) and associated 95% highest density intervals (HDIs) for the years 1990-2017. The observed number of calves born each year is shown in red. Expected calves born are calculated as the median number of females estimated to be adults and alive in each year who did not reproduce in the previous two years, from the model described in Pace et al. (2017). **(B)** The proportion of expected calves that were actually born. The values for each year are shown as black circles, and the average across all years (from 1990-2017, 27%) is shown as a red dashed line. A linear estimate of the trend over time is shown in blue, indicating a slight decrease over time (mean = -0.0034, SE = 0.0040).

## 4 DISCUSSION

The first major result from this work is the identification and quantification of the degree to which nulliparous females—and females who delay reproduction until much later than expected —are contributing to the reproductive problem (**Figure 1**). Indeed, by 2017 almost half (44%) of adult females had either not given birth yet or had just one calf; whereas just slightly over half (56%) of the adult females alive in that year had reproduced more than once and are therefore available for analyses of inter-birth intervals. These results, along with those from other work (e.g., Reed et al. 2022), show that non-reproductive females, and females who are delaying reproduction until much later in life, represent a major component of the reproductive problem, and should be considered along with data on inter-birth intervals when quantifying reproductive performance.

Second, our approach provides clear reproduction “targets” for each year, providing a scientific baseline against which actual reproductive performance can be compared. This approach shows that in none of the 28 years considered were the number of calves close to those expected (**Figure 2**). This result is alarming, and demonstrates the magnitude of the degree to which reproduction is compromised within this species.

Lastly, our approach provides a clear and scientific method to quantify and monitor the reproductive problem over time. This approach indicates that, on average, the species is reproducing at only about 27% of its expected rate, or put another way, the reproductive rate is about 73% lower than their known potential. Such information is critical for quantifying the relative degree to which reduced reproductive performance is limiting the recovery potential of the species. For example, future analyses could compare observed population trends with those expected if reproduction rates were not compromised, as well as those expected if anthropogenic mortalities were non-existent, to quantify the relative contribution of anthropogenic mortality and compromised reproduction on species recovery. Additionally, this quantification of the reproductive problem will aid research programs focused on identifying what factors are limiting reproductive performance and the degree to which they are doing so.

## ACKNOWLEDGMENTS

We thank all members of, and contributors to, the North Atlantic Right Whale Consortium for their dedication, continuing support, and data sharing policies and procedures. T. Frasier’s right whale research is supported by Fisheries and Oceans Canada (DFO), the Natural Sciences and Engineering Research Council of Canada (NSERC), Genome Atlantic, Genome Canada, and Research Nova Scotia.

